# Snord67 promotes lymph node metastasis and regulates U6-mediated alternative splicing in breast cancer

**DOI:** 10.1101/2020.08.28.272617

**Authors:** Yvonne L. Chao, Yinzhou Zhu, Hannah J. Wiedner, Yi-Hsuan Tsai, Lily Wilkinson, Alessandro Porrello, Amanda E.D. Van Swearingen, Lisa A. Carey, Jimena Giudice, Christopher L. Holley, Chad V. Pecot

## Abstract

Small nucleolar RNAs (snoRNAs) have long been considered “housekeeping genes”, important for ribosomal biogenesis and protein synthesis. However, there is increasing evidence that this largely ignored class of non-coding RNAs (ncRNAs) also have wide-ranging, non-canonical functions in diseases, including cancer. SnoRNAs have been shown to have both oncogenic and tumor suppressor roles, yet whether snoRNAs regulate metastasis is unknown. Here we show that expression of certain snoRNAs are enriched in lymph node (LN) metastases in a micro-surgical, immune-competent mouse model of breast cancer. We identify the snoRNA Snord67 as a key regulator of LN metastasis. Knockout of Snord67 resulted in significantly decreased LN tumor growth and subsequent development of distant metastases. This was associated with loss of targeted 2’-*O*-methylation on the small nuclear RNA U6, a component of the spliceosome. RNA sequencing revealed distinct alternative splicing patterns in Snord67 knockout cells. Using rapid autopsy breast cancer cases, we found that matched human primary tumor and LN metastases revealed similar alternatively spliced genes, including several that are known to contribute to cancer. These results demonstrate that Snord67 is critical for growth of LN metastases and subsequent spread to distant metastases, and suggest that snoRNA-guided modifications of the spliceosome represent a previously unappreciated, yet targetable pathway in cancer.

## Introduction

Distant metastases are the primary cause of cancer-related mortality. Metastasis to lymph nodes (LNs), in particular, correlates with both increased risk for distant metastases as well as poor prognosis (1). However, the unclear survival benefit from surgical resection of axillary LNs (AxLNs) in breast cancer suggests that while LN metastases are clinically important, the biologic relationships between LN metastases and distant metastases remain understudied (2). Historically, LN metastases have been considered a surrogate for the ability of cancer cells to metastasize via hematogenous dissemination; however, recent studies have demonstrated that LN metastases can directly give rise to distant metastases via lymphatic spread (3-5). Moreover, the LN harbors a unique microenvironment to which cancer cells must adapt. For example, tumor metastasis to LNs requires an adaptation to the fatty acid-rich microenvironment, and cancer cells accordingly up-regulate lipid metabolism pathways to survive (6). However, the mechanisms governing survival of cancers cells within LNs and spread to distant organs from LNs remain poorly understood.

To investigate the mechanisms that drive LN metastasis, we recently developed a micro-surgical mouse model using 4T1 breast cancer cells, which form tumors that histopathologically resemble triple-negative breast cancer, but are more similar to the luminal molecular subtype (7,8). Using this model, we demonstrated that micro-injected AxLN tumors are capable of spontaneously establishing distant lung metastases and that metastasis via the lymphatic route is more efficient than hematogenous dissemination. By evaluating protein-coding genes that were differentially expressed in micro-injected AxLNs, we found that cancer cells transiently upregulate the chromatin modifier histone deacetylase 11 (HDAC11) within the LN, yet decrease HDAC11 expression to escape the LN prior to establishing distant metastases(7). Given that non-coding RNAs (ncRNAs) provide layers of dynamic epigenetic regulation of gene expression, we investigated whether ncRNAs were also differentially expressed between LN metastases and primary tumors and distant metastases. ncRNAs are a broad class of RNAs that represent a significant component of the epigenetic landscape (9-13). Notably, microRNAs (miRs) and long ncRNAs (lncRNAs) are well established for regulating metastatic biology via extensive pre- and post-transcriptional regulation of cell fate (14,15).

## Results

### snoRNA expression is increased in lymph node metastases

Using the 4T1 model, we generated sub-clones of cancer cells obtained from mammary fat pad (MFP) tumors, microsurgically-injected AxLN tumors, and spontaneous lung metastases derived from AxLN tumors (AxLN-LuM). We then performed microarray profiling to identify ncRNAs that were differentially expressed in AxLN tumors compared to both MFP tumors and AxLN-LuM (Figure 1A and Supplemental Figure 1). Although numerous miRs and lncRNAs were in the broad classes of ncRNAs represented on the microarray, AxLN sub-clones were markedly enriched for small nucleolar RNAs (snoRNAs). Of the 30 ncRNAs that were differentially expressed in AxLN sub-clones compared with either the MFP or AxLN-LuM (total ncRNAs profiled, n=10,043), 67% were identified as snoRNAs; yet snoRNAs comprised only 12% of the total ncRNAs profiled (Figure 1B and Supplemental Table 1). SnoRNAs are small ncRNAs less than 300 nucleotides in length that primarily guide post-transcriptional modification of target RNAs via sequence complementarity at a region called the antisense element (Figure 1C). The two subtypes of snoRNAs, box C/D and box H/ACA, guide either 2’-*O*-methylation (box C/D) or pseudouridylation (H/ACA) of the target RNAs. The canonical targets of snoRNAs are ribosomal RNAs (rRNAs) and small nuclear RNAs (snRNAs), but snoRNA-guided modification of mRNA is also possible (16-20). In cancer, snoRNAs have been shown to have both oncogenic and tumor suppressor roles, yet whether and how snoRNAs regulate metastasis is unknown (21-24). The top six box C/D snoRNAs identified in the profiling analyses were validated using quantitative real-time polymerase chain reaction (qPCR) and demonstrated increased snoRNA expression in AxLN subclones compared to MFP and AxLN-LuM, with the most significant changes observed in Snord67 and Snord111 (Figure 1D). We generated matched pairs of MFP tumors and *de novo* LN metastases and then evaluated whether increased Snord67 and Snord111 expression was due to an artifact of microsurgical injection. For the majority of the matched pairs, expression of both snoRNAs was increased in the LN metastasis sub-clones compared to the MFP sub-clones, suggesting that the LN microenvironment drives transient expression of snoRNAs (Figure 1E).

**Figure 1.**
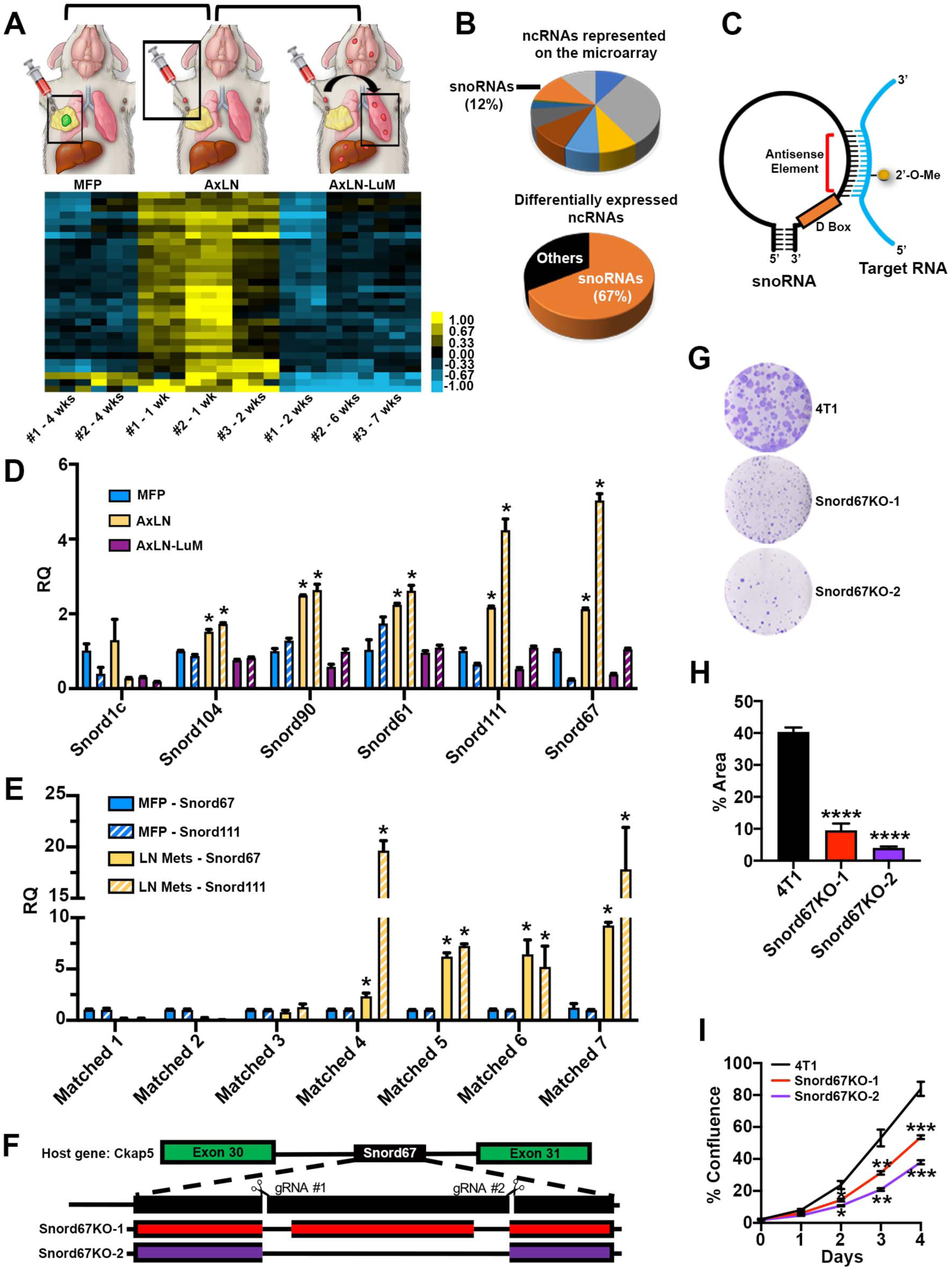
Expression of snoRNAs are increased in LNs. (**A**) Composite microarray showing ncRNAs that are up- or down-regulated in pairwise comparisons of: AxLN tumors (n=3) compared to MFP tumors (n=2), and AxLN tumors (n=3) compared to lung metastases derived from AxLN tumors (AxLN-LuM) (n=3). Pairwise comparisons are in brackets. Tumors were harvested at the indicated timepoints and then expanded *ex vivo* to generate subclones. Subclones were run in three biologic replicates on the microarray. (**B**) Pie chart of the categories of ncRNAs in the whole microarray (top) and within differentially expressed genes (bottom). (**C**) Schematic illustrating the structure of snoRNA and interaction with target RNA. (**D**) Relative quantification (RQ) of snoRNAs identified on the microarray in AxLN and AxLN-LuM subclones compared to MFP as evaluated by qPCR. Two subclones from each tumor site (MFP, AxLN, and AxLN-LuM) were assayed in technical triplicate, statistical significance calculated by ANOVA, *p<0.05. (**E**) qPCR RQ of Snord67 and Snord111 expression in de novo LN metastases compared to matched MFP tumors. Subclones were generated *ex vivo* in n=7 mice and run in technical triplicate, statistical significance calculated by ANOVA, *p<0.05. (**F**) Schematic showing the double nicking strategy for CRISPR-Cas9 gene deletion of snoRNAs. (**G-H**) Colony formation assay evaluating tumorigenesis. Cancer cells were plated in triplicate and then colonies were stained and quantified in relation to the area of each well that was covered. Statistical significance was calculated by ANOVA, ****p<0.0001. (**I**) Cell proliferation assay of 4T1 wild-type and Snord67 knockout (Snord67KO) cells. Proliferation was captured as time lapse images using Incucyte over 5 days. Growth was quantified as % confluence of each well. n=4 biologic replicates. Statistical significance was calculated by ANOVA, **p<0.01, ***p<0.001.

### Loss of Snord67 decreases proliferation and tumorigenesis

We next generated knockouts of Snord67 and Snord111 using the CRISPR/Cas9 system to evaluate the function of these snoRNAs within 4T1 breast cancer cells. Due to concerns that a single genomic deletion would not be sufficient to knockdown expression of snoRNAs (25), we employed a double nicking strategy by using paired guide RNAs to introduce reduce off-target mutagenesis (Figure 1F) (26). Two single-cell knockout clones for both Snord67 and Snord111 were isolated, and genomic deletions at the predicted sites were confirmed by DNA sequencing. The double nicking strategy resulted in clones with either two separate deletions or a larger deletion spanning the target sites. Loss of snoRNA expression in Snord67 and Snord111 knockout clones was verified by qPCR (Supplemental Figure 2A). snoRNAs are typically encoded within introns of host genes, and regulation of snoRNA expression can be dependent or independent of host gene transcription levels (27,28). Expression of host genes *Ckap5* (Snord67) and *Sf3b3* (Snord111) was not affected in knockouts (Supplemental Figure 2B-C). We then evaluated the functional significance of snoRNA depletion in breast cancer cells. Compared to 4T1 wild-type (WT) cells, loss of Snord67 resulted in significantly decreased colony forming ability and proliferation, while loss of Snord111 had no effect on either phenotype (Figure 1G-I and Supplemental Figure 2D-F). There was no effect on migration in either Snord67 or Snord111 knockouts (Supplemental Figure 2G). These results indicate that expression of Snord67, but not Snord111, is important for the growth and tumorigenicity of breast cancer cells *in vitro*.

### Snord67 is necessary for LN tumor growth and distant metastasis

To further evaluate the effect of Snord67 on LN tumor growth and distant metastasis, we micro-injected 4T1-mCherry/Renilla-luciferase-expressing wild-type or Snord67 knockout cells into the AxLN of immune-competent mice (Figure 2A). Mice that were injected with Snord67 knockout cells exhibited significantly decreased growth of AxLN tumors (Figure 2B). Using an assay to detect luciferase activity in cancer cells that had metastasized to the lungs, we found that mice injected with Snord67 knockout cells also developed significantly fewer distant lung metastases four weeks after AxLN injection (Figure 2C). To evaluate whether our observations on tumor growth were due to isolation of clonal populations of Snord67 knockout cells, we then tested the effects of abrupt inhibition of Snord67 expression using second generation antisense oligonucleotides (ASOs). The Snord67 ASO was designed to target the antisense element of Snord67 and was chemically-modified with flanking 2’–methoxyethyl (MOE) bases for enhanced stability, resistance to nuclease degradation, and avoidance of immunogenicity (Figure 2D-E). Compared with a control ASO, *in vitro* treatment of 4T1 cells with Snord67 ASO resulted in significantly decreased Snord67 expression in 4T1 cells (Supplemental Figure 2H). To test the effects of therapeutic inhibition of Snord67 expression *in vivo*, 4T1 WT cells were micro-injected into AxLNs to generate tumors. Two weeks after injection, mice were randomly distributed to the following treatment groups: 1) vehicle (PBS), 2) negative control ASO 48 mg/kg (mpk), 3) Snord67 ASO 12 mpk, 4) 24 mpk or 5) 48 mpk (Figure 2F). Baseline AxLN tumors were measured by caliper and daily subcutaneous treatments were administered for six consecutive days. AxLN tumors were measured and harvested 24 hours after the last treatment. Mice treated with 24 and 48 mpk of Snord67 ASO demonstrated significantly reduced tumor growth at 20 days post-LN injection, compared to vehicle treated mice (Figure 2G). Treatment with all tested doses of Snord67 ASO, but not PBS or negative control ASO, resulted in significant inhibition of Snord67 expression in AxLN tumors (Figure 2H). As inhibition of Snord67 expression but not inhibition of AxLN tumor growth was observed in the 12 mpk treatment group, it is possible that the 24 and 48 mpk doses of Snord67 ASO resulted in more rapid and sustained inhibition of Snord67 expression, and perhaps an anti-tumor effect could be detected with longer treatment of 12 mpk. These results demonstrate that ASOs can potently and specifically target snoRNAs in tumors, and may have therapeutic potential. Finally, to evaluate whether Snord67 may have effects on distant metastasis colonization once the cancer cells enter circulation, we utilized an experimental metastasis model. Consistent with reduced spontaneous lung metastases from LNs (Figure 2C), we observed a dramatic reduction of lung metastases in both Snord67 knockout groups compared with 4T1 WT cells (Figure 2I-J). Taken together, these results indicate that Snord67 is essential for both LN tumor growth and subsequent metastasis to distant sites.

**Figure 2.**
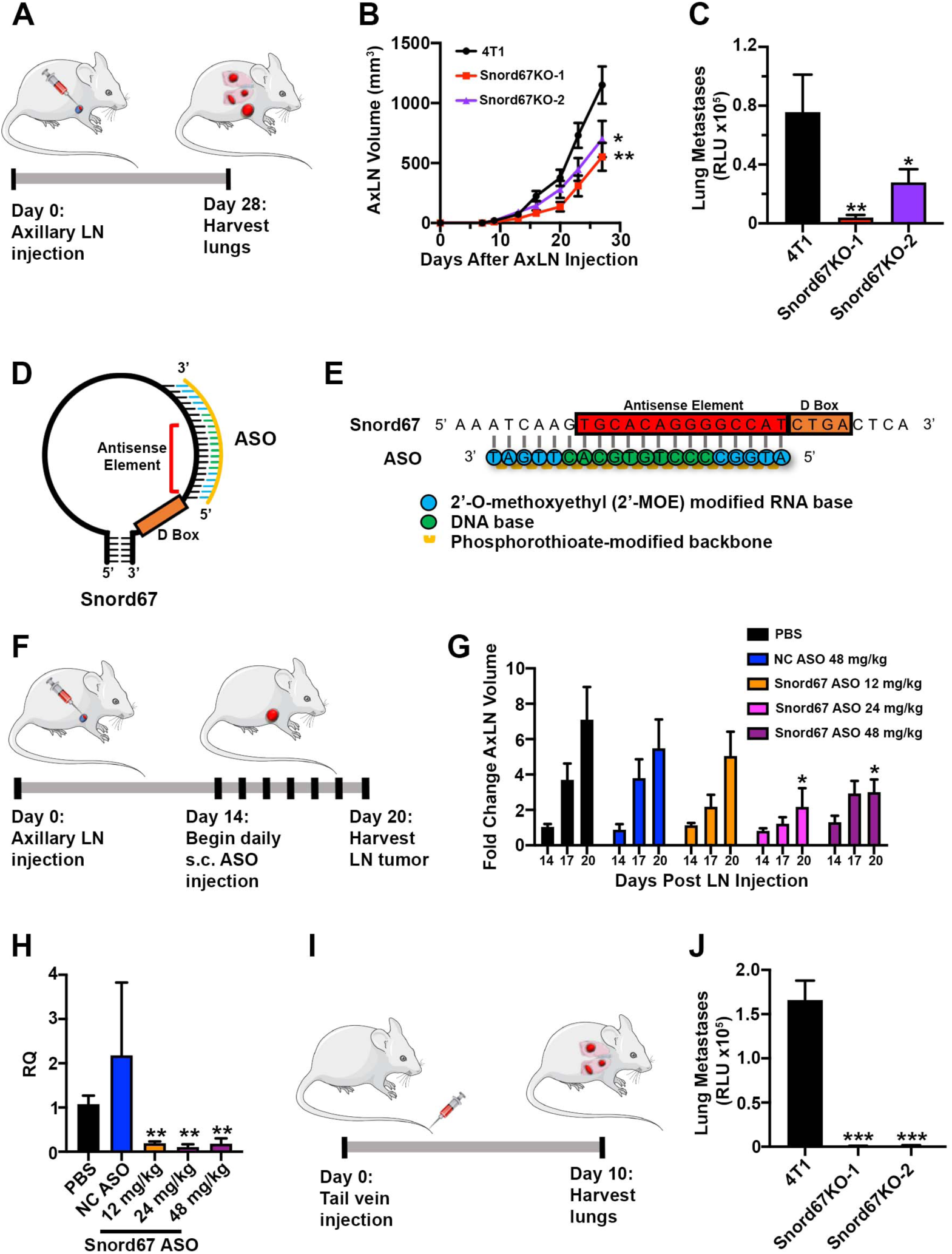
Snord67 is necessary for AxLN tumor growth and metastasis. (**A**) Schematic of microsurgical injection of AxLN leading to subsequent lung metastases. (**B**) AxLN tumor volumes as measured by calipers twice weekly (n=10 mice per group). Statistical significance was calculated by ANOVA, *p<0.05, **p<0.01. (**C**) Lungs were harvested at day 28 after AxLN injection and then digested into a single cell suspension. Relative Luciferase Units (RLU) were measured by luminometer to quantify lung metastases. Measurements were performed in technical triplicate. Statistical significance was calculated by ANOVA, *p<0.05, **p<0.01. (**D-E**) Design of ASO targeting the antisense element of Snord67. The ASO was designed with a phosphorothioate-modified backbone and 2’-methyoxyethyl-modified bases flanking the 5’ and 3’ ends. (**F**) Schematic of AxLN injection to form tumors, followed by daily treatment of ASO. AxLN tumors were harvested at day 20 following injection. (**G**) AxLN tumor volumes were measured by caliper measurements prior to ASO treatment, once during, and on day of AxLN tumor harvest. n=5 mice per treatment. Statistical significance was calculated by ANOVA, *p<0.05. (**H**) Snord67 expression by qPCR in AxLN tumors following treatment with ASO. RQ was calculated in comparison to the PBS-treated mice and expression of Snord67 was normalized to Gapdh and Rplp0. Dose of NC ASO was 48mg/kg. Statistical significance was calculated by ANOVA, **p<0.01. (**I**) Schematic of experimental metastasis experiment. Cancer cells were injected into tail veins and subsequently metastasized to lungs. (**J**) Ten days after tail vein injection, lungs were mechanically and enzymatically digested. Cells were then lysed and Relative Luciferase Units were measured in triplicate. Statistical significance was calculated by ANOVA, ***p<0.001.

### Snord67 knockouts exhibit distinct alternative splicing patterns

To elucidate the mechanisms of how Snord67 contributes to metastatic progression, we used Ribose Oxidation Sequencing (RibOxi-Seq) and RNA-Sequencing (RNA-Seq) to evaluate differential 2’-*O*-methylation and gene expression between 4T1 and Snord67 knockout cells. Snord67 is known to guide modification of U6 snRNA, so we first analyzed the 2’-*O*-methylation status of U6 using RibOxi-Seq (29). In this method, RNA is randomly fragmented and then oxidized such that fragments with ribose-methylated nucleotides at the 3’ end are protected while unmethylated nucleotides are degraded. Fragments ending in methylated nucleotides are positively selected and enriched for during library construction. Sites of 2’-*O*-methylation are then mapped using RNA-Seq and bioinformatics analysis. In 4T1 WT cells, all eight known sites of 2’-*O*-methylation on U6 were detected, but in Snord67 knockout cells there was a loss of methylated reads specifically at site C60, which is the site of Snord67-guided methylation (Figure 3A) (30). We next analyzed traditional RNA-Seq results for differential expression and found an unusually large number of differentially expressed genes (n=7,013 genes with adjusted p-value <0.05) (Supplemental Figure 3A). Because the known target of Snord67 is snRNA U6(30), we hypothesized that fundamental changes to splicing might therefore be the mechanism leading to widespread changes in gene expression and driving the phenotypes of the Snord67 knockouts. We therefore evaluated the RNA-Seq data for alternative splicing differences between 4T1 and Snord67 knockout cells (Figure 3B). Splicing was analyzed in terms of percent spliced in (PSI), or the ratio between reads including or excluding exons. mRNA isoforms with PSI values that changed greater than 10% (ΔPSI≥10) between 4T1 and Snord67 knockout cells were included in subsequent analyses. If site-specific methylation of U6 by Snord67 disrupted constitutive rather than alternative splicing, a substantial increase in retained intron (RI) events would have been expected. Instead, the differential AS events between 4T1 and Snord67 knockout cells were distributed among the different types of splicing, with most events categorized as skipped exon (SE) or mutually exclusive exons (MXE) (Figure 3C, Supplemental Figure 3B). We validated some of these AS events using reverse transcription PCR (RT-PCR) assays followed by gel electrophoresis, and quantification of PSI and showed strong correlation between the RNA-Seq and RT-PCR data (Figure 3D-F, Supplemental Figure 4).

**Figure 3.**
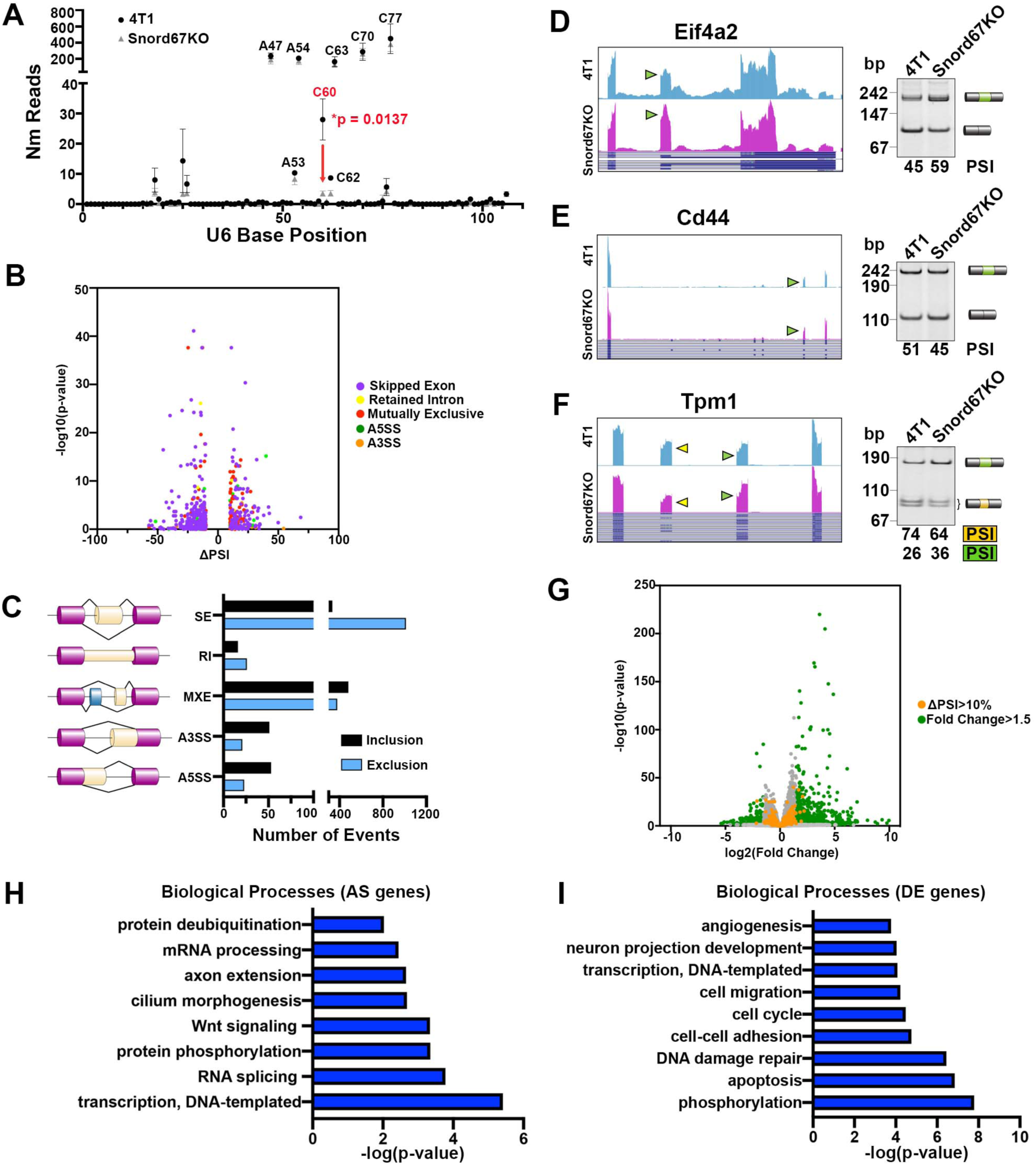
Knockout of Snord67 results in loss of 2’-O-Me and alterations in alternative splicing. (**A**) RibOxi-Seq mapping of methylation counts on snoRNA-directed U6 sites in 4T1 and Snord67 knockout (Snord67KO) cells. (**B**) Volcano plot of differential AS events with ΔPSI >10% and p-value <0.05. SE = skipped exon; RI = retained intron; MXE = mutually exclusive exons; A3SS = alternative 3’-splice site; A5SS = alternative 5’-splice site. (**C**) Distribution of AS events in Snord67KO cells. (**D-F**) RNA-Seq data displayed by the UCSC browser - mm10 (left) and RT-PCR assays (right) for genes with AS between 4T1 and Snord67KO cells. Arrows indicate alternatively spliced exons. bp, base pairs; PSI, percent spliced in. **(G**) Volcano plot of differentially expressed genes between 4T1 and Snord67KO cells, plotted as the log2(Fold Change) on the x-axis and the -log10(p-value) on the y-axis. Genes that are differentially expressed with an absolute log2(fold change) greater than 1.5 are green. Genes that were found to be differentially spliced with a ΔPSI>10% are yellow. (**H-I**) Gene ontology analysis of genes that are differentially alternatively spliced (H) or differentially expressed (I) between 4T1 and Snord67KO cells using the DAVID functional annotation tool. Terms with FDR<0.05 were plotted according to p-value.

### Snord67 knockout cells demonstrate disparate differential gene expression and AS patterns

There was little overlap between genes that were differentially alternatively spliced and those that were differentially expressed (Figure 3G). Gene ontology analysis of the alternatively spliced genes showed categorical enrichment for transcription, RNA splicing, and mRNA processing (Figure 3H). However, genes that were differentially expressed between Snord67 knockouts and 4T1 wild-type were largely categorized within biological processes such as apoptosis, DNA damage repair, cell cycle, and cell adhesion (Figure 3I). To evaluate whether Snord67-directed methylation of each mRNA could be contributing to AS independent of U6 modification, we used RibOxi-Seq to evaluate whether there was disrupted methylation of pre-mRNA of the validated AS genes. Methylation mapping of mRNA demonstrated no significant difference in 2’-*O-*methylation patterns between 4T1 and Snord67 knockout cells in regions surrounding the AS event (Supplemental Figure 5A-E). Taken together, these results led us to two conclusions. First, Snord67 broadly impacts transcript abundance and alternative splicing in non-overlapping genes, and thus, non-overlapping cellular functions. Second, Snord67 regulates the splicing of genes that themselves are involved in splicing and mRNA processing. These results highlight that evaluating only quantitative changes in transcript abundance may ignore a considerable degree of isoform diversity that could contribute to cancer progression.

### LN metastases in humans with breast cancer exhibit similar alternative splicing changes as Snord67 knockouts

Using a cohort of six metastatic breast cancer patients obtained through the University of North Carolina Breast Cancer Rapid Autopsy Program (UNC RAP) (31), we examined AS differences between matched primary breast tumors and LN metastases (Figure 4A). AS differences were observed between matched LN metastases and primary tumors in all patients and distributed among event types (Supplemental Figure 6A-F, left panels). Functional annotation of differential AS between primary tumors and LN metastases, revealed that all six patients exhibited AS events in genes involved in RNA splicing, similar to what we observed in the Snord67 knockout cells, as well as cell-cell adhesion (Supplemental Figure 6A-F, right panels). We identified a set of 60 genes with differential AS events between LN metastases and primary tumors that were observed in all six RAP human samples (Figure 4B, left). We evaluated for overlap between the 60 UNC RAP genes and the AS genes discovered in Snord67 knockouts compared to 4T1 WT cells, surmising that these would be most likely to be clinically relevant, and found nine genes in common (Figure 4B, right). AS events for these genes common to both UNC RAP samples and Snord67 knockout cells were frequently conserved between mouse and human (Figure 4C-E). Because small ncRNAs (including snoRNAs) were excluded during RNA isolation and library preparation for RNA-Seq, we were unable to evaluate Snord67 expression in the UNC RAP samples. Interestingly, several of the AS events common to both analyses have been previously shown to contribute to cancer progression. In particular, AS of *CD44, TPM1*, and *EIFf4A2* genes have been demonstrated to have cancer-specific and even metastasis-specific AS isoforms (8,32-36). Taken together, these results support a model in which alternative splicing events are differentially regulated in primary tumors and LN metastases, both in breast cancer patients and our animal model.

**Figure 4.**
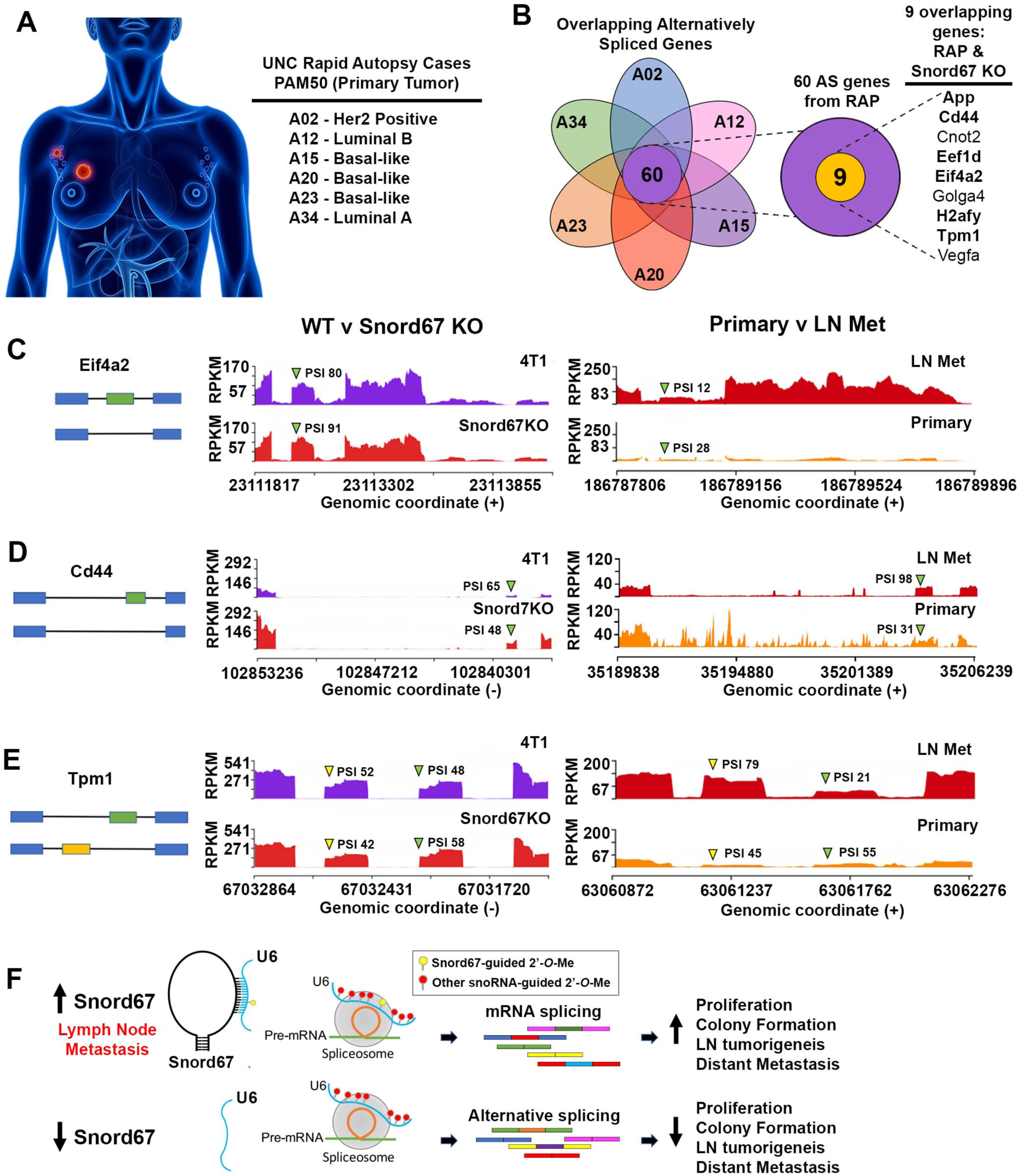
Validation of clinically relevant AS genes using rapid autopsy samples. (**A**) Graphic representation of matched primary breast tumors and LN metastases obtained from patients through the UNC Rapid Autopsy Program (RAP), which are annotated as A02, A12, A15, A20, A23, and A34. The PAM50 molecular subtypes of each case are listed. (**B**) Venn diagram showing the overlap in 60 AS genes between RAP samples. Out of these 60 genes, nine were also differentially spliced between 4T1 WT and Snord67KO cells (dotted inset and listed on right). Bolded genes exhibited conserved AS events between human (RAP) and mouse (Snord67KO) samples. (**C-E**) Sashimi plot visual representation from MISO showing AS events that were found to be differentially spliced between 4T1 WT and Snord67KO cells (left plot) and between LN metastases and primary tumors in UNC RAP patient samples (right plot). Sashimi plots are showing the splice junctions aligned on the genomic coordinates of each gene. Arrows indicate the alternative exon and the corresponding PSI. RPKM = reads per kilobase of transcript. (**F**) Proposed model depicting how Snord67-guided methylation of U6 leads to changes in splicing programs which might promote LN tumor growth and distant metastasis.

## Discussion

Metastasis is a highly complex, multi-step cascade that requires tremendous phenotypic plasticity in cancer cells in order to survive multiple microenvironments (37). Moreover, our previous work showed that metastasis via the lymphatic route was more efficient compared to hematogenous dissemination (7). In combination with our prior study showing that chromatin modifier HDAC11 is dynamically regulated within LN, the findings presented here underscore that distinct mechanisms within the unique LN microenvironment can promote metastatic tumor growth. Here, we show that expression of key snoRNAs are upregulated in the LN. We identified Snord67 as a novel ncRNA with essential roles for growth and survival in LNs, and perhaps even more importantly, for subsequent metastasis to distant sites. A previous study showed that loss of a specific box H/ACA snoRNA, SNORA23, led to decreased tumor growth and metastasis from a xenograft model of pancreatic cancer; however, the mechanisms of how snoRNAs regulate metastasis have not been delineated (38). Commonly considered to be “housekeeping genes”, several novel functions of snoRNAs have recently been identified, such as miRNA-like gene repression and mRNA 3’ processing (39-41). Our results suggest a mechanism in which site-directed methylation of U6 by Snord67 is required for the baseline splicing patterns, and loss of this modification leads to a new equilibrium of alternative splicing (Figure 4F). These altered splicing patterns are also observed in a cohort of metastatic breast cancer patients. Whether these altered splicing patterns are associated with dysregulated Snord67 expression in patients and whether the splice variants themselves have specific roles in cancer progression remains to be determined. Besides epigenetic regulation, AS is another mechanism by which cancer cells transiently and dynamically regulate genes that enable metastasis (42-44). Two studies recently described a role for LARP7, a RNA-binding protein that mediates the interaction between U6 and U6-targeting snoRNAs like Snord67 (45,46). Knockout of LARP7 in human embryonic kidney cells resulted in loss of methylation at all known U6 sites, as well as resultant changes in AS. How snoRNA-directed modifications on U6 leads to changes in AS is unknown. It is possible that 2’-*O*-methylation of U6 affects binding of splicing factors or RNA-binding proteins, or perhaps leads to conformational changes in U6 that alter association with other spliceosome components (17,47,48). Finally, our results reveal that ASO technology may be useful for therapeutically targeting cancer metastasis, through either the use of ASOs to target dysregulated, pro-metastatic snoRNAs, or potentially through the use of ASOs that are specifically designed to target metastasis-specific AS events. For example, splice-site-specific ASOs such as nusinersen have demonstrated remarkable clinical efficacy in other applications like spinal muscular atrophy (49,50). Future work directed at further elucidating the mechanisms of how snoRNAs promote metastasis through alternative splicing could therefore lead to novel strategies for treating cancer patients.

## Methods

### Cell lines

4T1 cells were obtained from the ATCC and maintained in RPMI containing 10% FBS. 4T1 cells were transduced with lentiviral constructs expressing either GFP/firefly luciferase or mCherry/renilla luciferase and then selected and maintained in puromycin (4 μg/ml). All cells were routinely tested for mycoplasma using a Lonza MycoAlert Detection kit (LT07-418).

### Constructs and key reagents

All-in-one CRISPR-Cas9 clones targeting Snord67 were purchased from Genecopoeia (vector pCRISPR-CG02). sgRNA sequences were designed with help from CRISPOR (51) and sequences are listed in the table below. Cells were transiently transfected with pairs of Cas9-sgRNA constructs by electroporation using the Neon Transfection System (Invitrogen). Single cell clones were generated using serial dilution. To confirm Snord67 gene deletion, DNA was isolated using the DNeasy Blood and Tissue Kit (Qiagen, Cat 69504) and then sequenced (Eton Biosciences). Sequencing primers are provided in Supplemental Table 2. Negative control and Snord67 ASOs were synthesized by IDT.

### qPCR

RNA was purified from cells using the ZymoQuick RNA miniprep kit (Zymo Research, R1057). cDNA was synthesized using the SuperScript First-Strand cDNA Synthesis Kit (Thermo Fisher, 11904018) using gene-specific primers for stem-loop qPCR or the iScript cDNA Synthesis Kit (BioRad, 1708890). cDNA was analyzed using SYBR Green Master Mix (BioRad, 1525271). PCR was run on a StepOnePlus qPCR machine (Applied Biosystems). Data was analyzed using the ΔΔCt method, and experiments were normalized to Gapdh and Rplp0.

### Migration Assay

A total of 25,000 4T1 cells were added in serum-free medium to BioCoat Control Cell Culture Inserts with 8-μm pores (Corning, 354578) that were pre-coated with 10 μg/ml type 1 rat tail collagen on the bottom of the inserts. RPMI medium containing 10% FBS was added to the lower chambers as the chemoattractant. After 18 hours, cells that had migrated to the bottom of the filter were fixed and stained using the Protocol Hema 3 staining kit (Fisher Scientific, 22122911). Membranes were mounted onto glass slides, and images were taken using a Nikon microscope. Migrated cells were enumerated using CellProfiler open source image analysis software.

### Colony formation assay

After treatment, cells were trypsinized, counted, and 1,000 or 5,000 cells were plated in triplicate in six-well plates containing complete RPMI medium and drug, as indicated. Cells were allowed to grow under standard conditions for at least 4 days until colonies were observed. For staining, 1 ml of crystal violet stain (0.05% crystal violet, 1% formalin, 1% methanol in PBS) was added to the cells. Images were taken using an Epson office scanner under film settings. Colonies were quantified using the ColonyArea plugin on ImageJ (52).

### Proliferation Assay

Cells were plated at a density of 2,500 cells per well in a 96-well plate in 120 µl of RPMI containing 10% fetal bovine serum. The plate was placed in an IncuCyte ZOOM Live-Cell Imaging system (Essen Bioscience) (10x objective) and the percent confluence was recorded every two hours by both phase contrast and fluorescence scanning for 96 hours at 37 °C and 5% CO2. Images were analyzed using the Incucyte ZOOM software and the percentages of cell confluence were calculated over time.

### Mammary fat pad injection

For all *in vivo* studies, adult female Balb/c mice (6-8 weeks) were purchased from Taconic Farms. Cells were trypsinized and suspended in Matrigel at a 1:1 ratio, and 5,000 cells were injected directly into the 8^th^ mammary fat pad. Caliper measurements of subcutaneous tumor growth were taken twice weekly and the tumor volume was calculated as L X W^2^ where L is the greatest cross-sectional length across the tumor and W is the length perpendicular to L.

### Axillary lymph node injection

Mice were anesthetized, depilated, and subjected to surgical implantation of 5,000 (4T1) cells in a total volume of 1 μl Hank’s Buffered Saline Solution (HBSS). Injections were performed using a dissecting microscope and a 10-μl Hamilton syringe and custom-made microtip Pasteur pipette. Caliper measurements of tumor growth were taken twice weekly, and the tumor volume was calculated as above.

### Tail vein injection

Mice were injected with 1×10^5^ cells in HBSS by tail vein, after which mice were monitored daily for health. Mice were sacrificed and analyzed at the indicated time point post-injection.

### ASO treatment

Second generation ASOs designed to target the antisense element of Snord67 were designed with a phosphorothioate backbone with 10 central DNA bases flanked by 5 methoxyethyl (MOE) modified bases at the 5’ and 3’ ends and synthesized by IDT. Mice were micro-injected with 4T1 cancer cells in the AxLN as described above. Two weeks after injection of cancer cells into the axillary lymph node, PBS or ASO treatment was initiated. ASO was resuspended in 200ul PBS and injected subcutaneously in the scapular area daily for 6 total treatments.

### Ex vivo cell line establishment

Tumors were excised in a sterile fashion and minced in digestion medium. Tissue was then digested for 1 hour in 0.125% collagenase II, 0.1% hyaluronidase, 15 U/ml DNase, and 2.5 U/ml Dispase. Cells were then pelleted, subjected to ACK red blood cell lysis, and plated in 10-cm dishes containing complete RPMI medium and antibiotics as appropriate. For passaging and subsequent culture, *ex vivo* 4T1 sub-clones were selected with 6-thioguanine for several days until pure colonies were observed.

### Quantification of lung metastases

Lungs were extracted and placed into dissociation buffer that was composed of 1mg/ml collagenase type 2 (Worthington, #LS004177), 0.25U/ml neutral protease (Worthington, #LS02104), 4ug/ml Deoxyribonuclease (Worthington, #LS006343), and 500U/ml hyaluronidase (MP Biomedicals, #ICN10074090) resuspended in low-glucose DMEM (Gibco). Tissues were then mechanically minced using the gentleMACS™ Octo Dissociator (Miltenyi Biotec) and then enzymatically digested at 37 °C for 1 hour. Red blood cells were lysed with ACK Lysis buffer and then cells were plated following several washes with PBS. Cells were cultured at 37 °C for 1 week in complete growth media supplemented with 6-thioguanine. Renilla luciferase activity was then detected by the Renilla Luciferase Assay System (Promega, E2810) using a luminometer.

### Validation of alternative splicing by RT-PCR

Reverse transcriptase polymerase chain reaction (RT-PCR) was performed using the High Capacity cDNA Reverse Transcription Kit (Applied Biosystems) according to manufacturer instructions. PCR reactions were performed using GoTaq DNA Polymerase (Promega) following manufacturer protocols. Primers were designed to anneal to the constitutive exons flanking the alternative spliced exon based on RNA-seq data with Primer3 software. Primer sequences for splicing validation are shown above. PCR amplification was performed using a thermocycler (Eppendorf Mastercycler Ep Gradient) and with the following steps (i) 95 °C for 75 s, (ii) 27 cycles of 95 °C for 45 s, 57 °C for 45 s and 72 °C for 1 min, (iii) 72 °C for 1 min and (iv) 25 °C hold. Tpm1 PCR products were digested with PvuII at 37 °C for 1 h. Products were resolved via 6% PAGE and imaged using a ChemiDoc XRS (Bio-Rad). The percent spliced in (PSI) was estimated by densitometry (Chemidoc Image Lab Software, Bio-Rad) using a pUC19 DNA ladder calibration curve and according to the following equation: PSI = 100 x (Inclusion band)/(Inclusion band + Skipping Band).

### Microarray analyses

These microarray results are based on samples belonging to a larger experiment that we performed; some findings coming from this experiment have been previously published(7). Their annotation and raw files have been deposited in the Gene Expression Omnibus (GEO) repository, with the following accession code: GSE136031. Because those samples have already been described in that article and in its GEO metafile, this paragraph of bioinformatics methods aims to 1) summarize what can be found in the aforementioned publication and is relevant also for this new research 2) highlight differences in the data processing and analysis between that and this study and 3) document how the three heat maps that are shown in the main text and supplementary data were generated.

The only samples that are described in the GEO metafile of GSE136031 and were used for the data processing and analyses performed here are: 1) two baseline sample types (4T1-GFP-fLuc and 4T1-mCherry-rLuc), each as a quadruplicate, for a total of 8 samples 2) two mammary fat pad (MFP) sample types, injected in and harvested from MFP, each as a triplicate (6 samples) 3) three axillary lymph node (AxLN) sample types, injected in and harvested from AxLN, each as a triplicate (9 samples) 4) three axillary lymph node (-derived) lung metastases (AxLNLuM), injected in AxLN and harvested from lung metastases (LuM), each as a triplicate (9 samples). Samples were obtained at different time points (see the previously mentioned GEO metafile). The generation and normalization of the RNA expression values relied on the robust multi-array algorithm (RMA) (53,54). The expression measures of these two plates were combined through a joint scaling procedure (all 41,345 Affymetrix probes combined). However, for this research, we analyzed only those probes that were not included in our previous analysis, which had focused on protein-coding genes, and did not have a missing mRNA description or were not poorly characterized in the Affymetrix annotation. The analyzed samples are the same, with the important and already stated difference that the two studies focus on two different subsets of Affymetrix probes.

The analysis started with a tailored differential gene expression (vs. the two baselines) including genes: i) having a mean expression value for at least one sample triplet (i.e., a sample type) ≥ 50% or ≤ 50% than the mean in the two baselines ii) ≥ 25th percentile in terms of range (across the samples), calculated on the sub-matrix whose columns are the baselines and that sample triplet, and iii) for which every comparison between the three samples of that triplet and the baseline replicates followed either the mathematical relation of being > or of being <. Thereafter, different sample types were compared, following this scheme: a) MFP vs. AxLN, b) MFP vs. AxLNLuM, and c) AxLN vs. AxLNLuM. These three comparisons had to fulfill the requirement defined in the above point iii). For the purposes of this research, the two direct comparisons that are shown are MFP vs. AxLN as well as AxLN vs. AxLNLuM. For an overview of how the probes for the heat map that displays MFP vs. AxLN vs. AxLNLuM (24 samples total) were assembled, refer to Leslie et al, since the method used is the same (7).

Finally, we hierarchically clustered the matrix rows (each containing the gene expression values of an Affymetrix ID across the selected samples) on data that were log2-trasformed and mean-subtracted using publicly available software (55,56).

### RibOxi- and RNA-Sequencing and Analysis

Total RNA was isolated using the Purelink RNA Mini Kit (Invitrogen). Three biologic replicates were used for each sample. RNA samples were assessed using bioanalyzer to ensure sample quality. mRNA was then enriched using NEBNext® Poly(A) mRNA Magnetic Isolation Module. Enrichment was assessed by examining rRNA depletion using a bioanalyzer. For mRNA-seq library prep, the Kapa Stranded mRNA-Seq kit (Roche) was used. The methods for the RibOxi-Seq library prep were modified from a previously published protocol (29). In brief, Poly(A) enriched RNA was denatured for 3 min at 90°C and fragmented by Benzonase (final RNA concentration: 100 ng/µL, enzyme 0.5 units/100µL reaction) on ice for 90 minutes. Following extraction with acidic phenol-chloroform, the fragmented samples underwent 2 rounds oxidation-elimination cycles. The oxidized RNA was then ligated to the NEB Universal miRNA cloning linker using T4 RNA ligase 2, truncated KQ (NE Biolabs, M0373S) at 16°C overnight. 25uM RibOxi RT primer was then annealed to the reaction before ligation of 5’ RNA linker using T4 RNA ligase I at room temperature for 1 hour and subsequent reverse-transcription using Protoscript II (NE Biolabs, M0368S). cDNA was then purified with AmpureXP beads with 1:1.8 ratio. 10uM Illumina compatible forward and reverse PCR primers were added together with Q5 master mix (NE Biolabs, M0544S) followed by Ampure XP purification with 1:1 ratio. A custom cycling program was used to amplify the final PE libraries where 5 cycles were with annealing temperature of 55 °C, while the remaining 23 cycles had annealing temperature of 62 °C. Library QC was done with Agilent bioanalyzer 2200 with DNA1000 chips and reagents. Samples were then sequenced by the UNC High-throughput Sequencing Facility using a HiSeq 4000 (Illumina). Sequences of primers and oligos used in RNA-Seq and RibOxi-Seq are provided in Supplemental Table 3.

For mRNA differential expression analysis, RNA-sequencing data was mapped with *STAR* (2.6.1) using a mapping index of mm10. Extra adapter read-through sequences and short/empty insert reads were removed entirely. The resulting alignment files were processed with *featureCounts* to generate count matrices. Log-2-fold changes and FDR values were then obtained by importing the count matrix to R and perform analysis using *DESeq2* package.

For differential splicing analysis, the above mapping step-generated junction files were concatenated to a single junction file and was provided to *STAR* per instructions of *STAR* 2-pass mapping mode. The reads were then re-mapped to the newly generated index. The resulting BAM files were used as inputs for *rMATs* (4.0.2) and the resulting output was filtered with ΔPSI of 15%. For RibOxi-Sequencing analysis, *cutadapt* was used to remove read-through adapters since insert sizes are predicted to be small. *pear* was then used to merge each pair of reads into a single read with default parameters. The actual 3’-linker sequence was removed using *cutadapt*. Mapping was done against the index generated in the mRNA Seq analysis step or U6 snRNA/rRNA transcript sequences. The read 3’-end counts are then generated by processing BED files converted from alignment BAMs. Gene ontology analysis was performed using the DAVID Functional Annotation Tool (57).

### Alternative splicing analysis of UNC RAP samples

We analyzed a cohort of six breast cancer patients from the University of North Carolina Breast Cancer Rapid Autopsy Program (UNC RAP) for whom primary tumors (T), LN metastasis (L), and distant metastases (D) were collected as previously described(7,31). For all primary tumors and LN metastases with available RNAseq, we estimated the expression of alternative splicing Psi (Ψ value, Percent Spliced In) at event level using MISO exon-centric analysis(58). We created a MISO annotation file using the gencode release 22 (in GRCh38), and chose the “commonshortest” flanking rule while converting the gencode annotation to the splicing events using the rnaseqlib package (58). We then performed pairwise comparisons between primary tumor and LN metastasis within each of the six patients using the compare_miso function from MISO package. Any splicing changes with Bayes factor larger than 20 were considered as significant events. We then further checked if any of those significant events from human RAP samples were also observed in the mouse models. Because of the difficulty of converting human splicing events to mouse splicing events, we only consider the overlap at the gene level.

### Data Availability

The microarray data that support the findings of this study have been deposited in the Gene Expression Omnibus (GEO) data bank, accession code GSE136031.

### Statistics

For *in vivo experiments*, between 5 and 15 mice were assigned per treatment group; this sample size gave approximately 80% power to detect a 50% change in tumor weight with 95% confidence. Results *in* vitro and *in vivo experiments* were compared using Student *t* test (for comparisons of two groups) and analysis of variance (for multiple group comparisons). For values that were not normally distributed (as determined by the Kolmogorov-Smirnov test), the Mann– Whitney rank sum test was used. A *P* value less than 0.05 was deemed statistically significant. All other statistical tests for *in vitro* experiments were performed using GraphPad Prism 8 (GraphPad Software, Inc., San Diego, CA). The multiple hypothesis testing correction of these results was made using the FDR.

### Study approval

All animals were cared for according to guidelines set forth by the American Association for Accreditation of Laboratory Animal Care and the U.S. Public Health Service policy on Human Care and Use of Laboratory Animals. All mouse studies were approved and supervised by the University of North Carolina at Chapel Hill Institutional Animal Care and Use Committee.

## Author contributions

Conceptualization: Y.L.C. and C.V.P., Y.L.C., C.L.H, J.G., and C.V.P. designed the experiments, Y.L.C, H.J.W., L.W., A.E.D.V., and C.V.P. performed the experiments. Y.Z. performed RNA-Seq and RibOxi-Seq analysis. A.P. performed microarray data analysis. Y.T. performed RNA-Seq alternative splicing analysis on UNC RAP samples. L.A.C procured funding, resources, and patients for UNC RAP. Y.L.C. wrote the paper with input from all authors.

## Competing interests

The authors declare no competing interests.

## Acknowledgments

The authors acknowledge members of the Pecot lab for helpful discussions and feedback. We would also like to thank the patients and their families who generously contributed to the UNC Rapid Autopsy Program.

## Funding

The UNC High-Throughput Sequencing Facility is supported in part by an NCI Center Core Support Grant (CA016086) to the UNC Lineberger Comprehensive Cancer Center and the University Cancer Research Fund. The UNC Breast Cancer Rapid Autopsy Program is supported by a Susan G. Komen Scholar Award (L.A.C.) and the UNC Specialized Program of Research Excellence (SPORE) P50-CA58223. C.L.H and Y.Z. were supported by NIH R01HL146381, R01 GM135383, Duke Strong Start Physician-Scientist Award, and the Mandel Foundation. J.G. was supported by start-up funds from the University of North Carolina at Chapel Hill, the National Institutes of Health (NIH R01GM130866), and a Career Development Award from the American Heart Association (19CDA34660248). H.J.W. was supported in part by the NIH-NIGMS training award T32 GM119999 and by the NSF Graduate Research Fellowship Program (DGE-1650116). Y.L.C was supported by the John Pope Fellowship Award from UNC. C.V.P. was supported in part by the National Institutes of Health R01CA215075, a Mentored Research Scholar Grants in Applied and Clinical Research (MRSG-14-222-01-RMC) from the American Cancer Society, the Jimmy V Foundation Scholar award, the UCRF Innovator Award, the Stuart Scott V Foundation/Lung Cancer Initiative Award for Clinical Research, the University Cancer Research Fund, the Lung Cancer Research Foundation, the Free to Breathe Metastasis Research Award, the Susan G. Komen Career Catalyst Award and a NCBC translational research grant.

## Notes

**Disclosure of Potential Conflicts of Interest:** The authors disclose no potential conflicts of interest.

### Competing Interest Statement

The authors have declared no competing interest.

