## Supplemental Data for "Snord67 promotes lymph node metastasis and regulates U6-mediated alternative splicing in breast cancer"

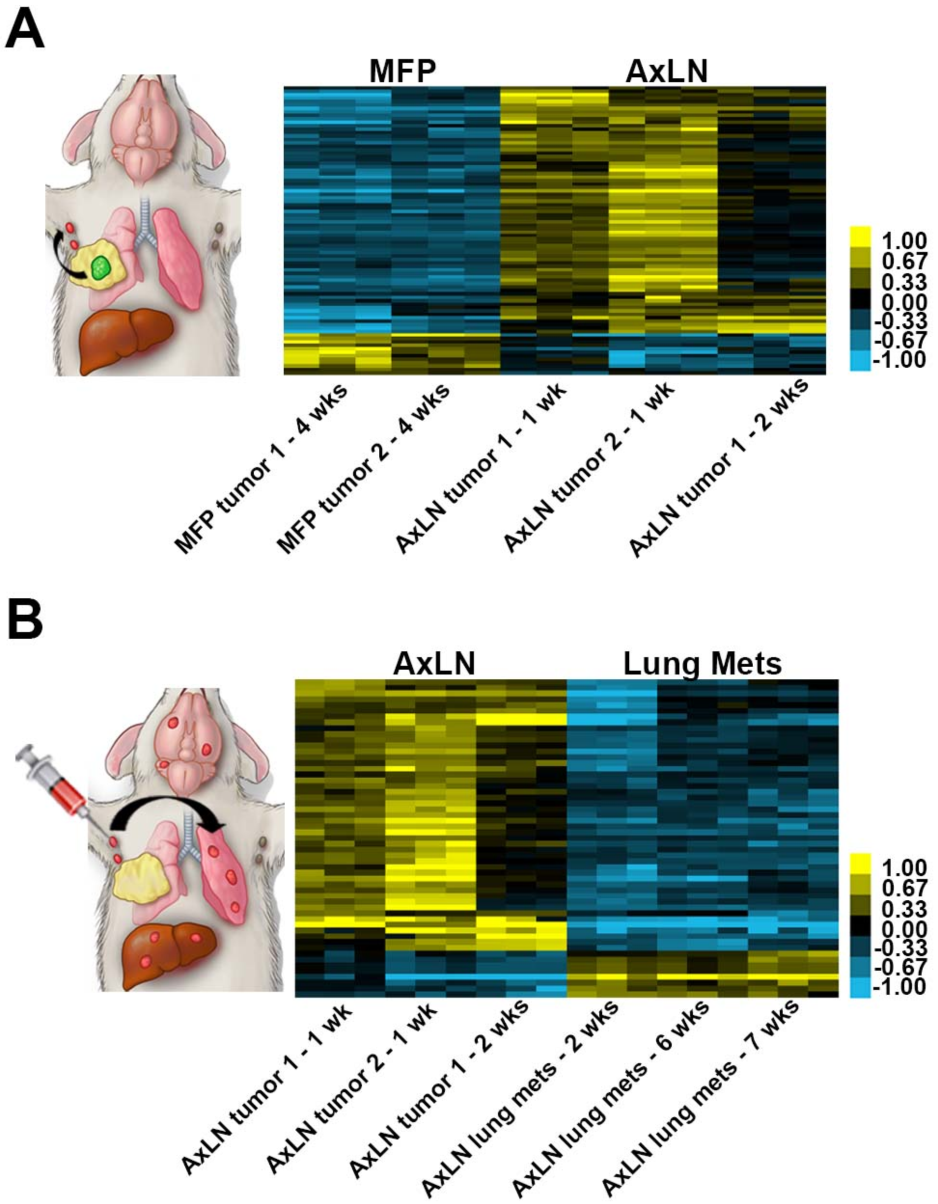


**Supplemental Figure 1. Non-coding RNAs are differentially expressed between mammary fat pad (MFP), axillary LN, and lung metastases.** (**A**) Pairwise comparison of differentially expressed ncRNAs in AxLN tumors (n=3) vs. MFP tumors (n=2). (**B**) Pairwise comparison of differentially expressed ncRNAs in AxLN tumors (n=3) vs. lung metastases derived from AxLN tumors (n=3). Tumors were harvested at the indicated timepoints and then expanded *ex vivo* to generate subclones. Subclones were run in three biologic replicates on the microarray.

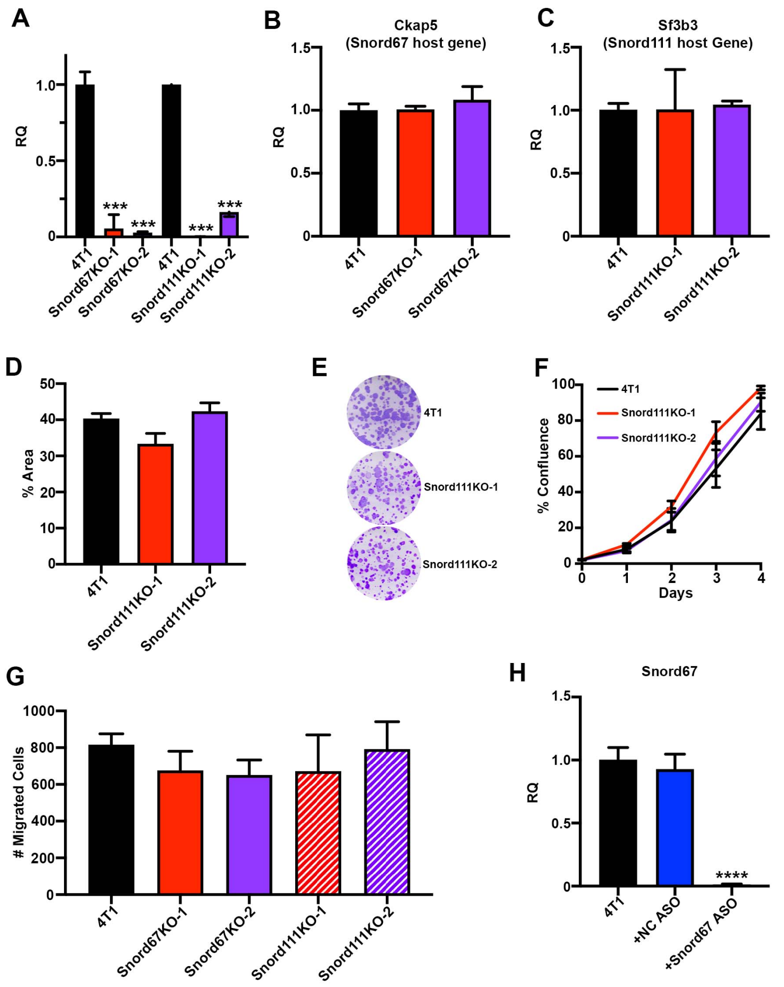


**Supplemental Figure 2. Snord67 knockout does not affect host gene expression or migration. (A)** Expression of Snord67 and Snord111 in 4T1 and CRISPR knockouts as quantified by qPCR and presented as relative quantification (RQ) compared to 4T1 WT. Samples were run in technical triplicate. Statistical significance was calculated by ANOVA, ***$p<0.001$. (**B-C**) Expression of Snord67 host gene CKAP5 and Snord111 host gene SF3B3 as quantified by qPCR and presented as RQ compared to 4T1 WT. Samples were run in technical triplicate. Statistical significance was calculated by ANOVA. (**D-E**) Colony formation assay demonstrating tumorigenesis. 4T1 and Snord111KO cancer cells were plated in triplicate and then colonies were stained and quantified after 7 days. Colonies were quantified in relation to the area of each well that was covered. Statistical significance was calculated by ANOVA. (**F**) Cell proliferation assay of 4T1 and Snord111KO. Growth of cells was captured as time lapse images using Incucyte over 5 days. Growth was quantified as % confluence of each well. N=4 biologic replicates. Statistical significance was calculated by ANOVA. (**G**) Quantification of migration of 4T1, Snord67KO, and Snord111KO cells by trans-well assay. Migration was measured as the number of cells on the underside of the migration chamber membrane after 18 hours. Statistical significance was determined by ANOVA. (**H**) qPCR relative quantification of Snord67 expression following 1uM ASO treatment for 48 hours *in vitro* compared to 4T1 untreated and treated with 1uM negative control (NC) ASO. Statistical significance was calculated by ANOVA, ****$p<0.0001$.

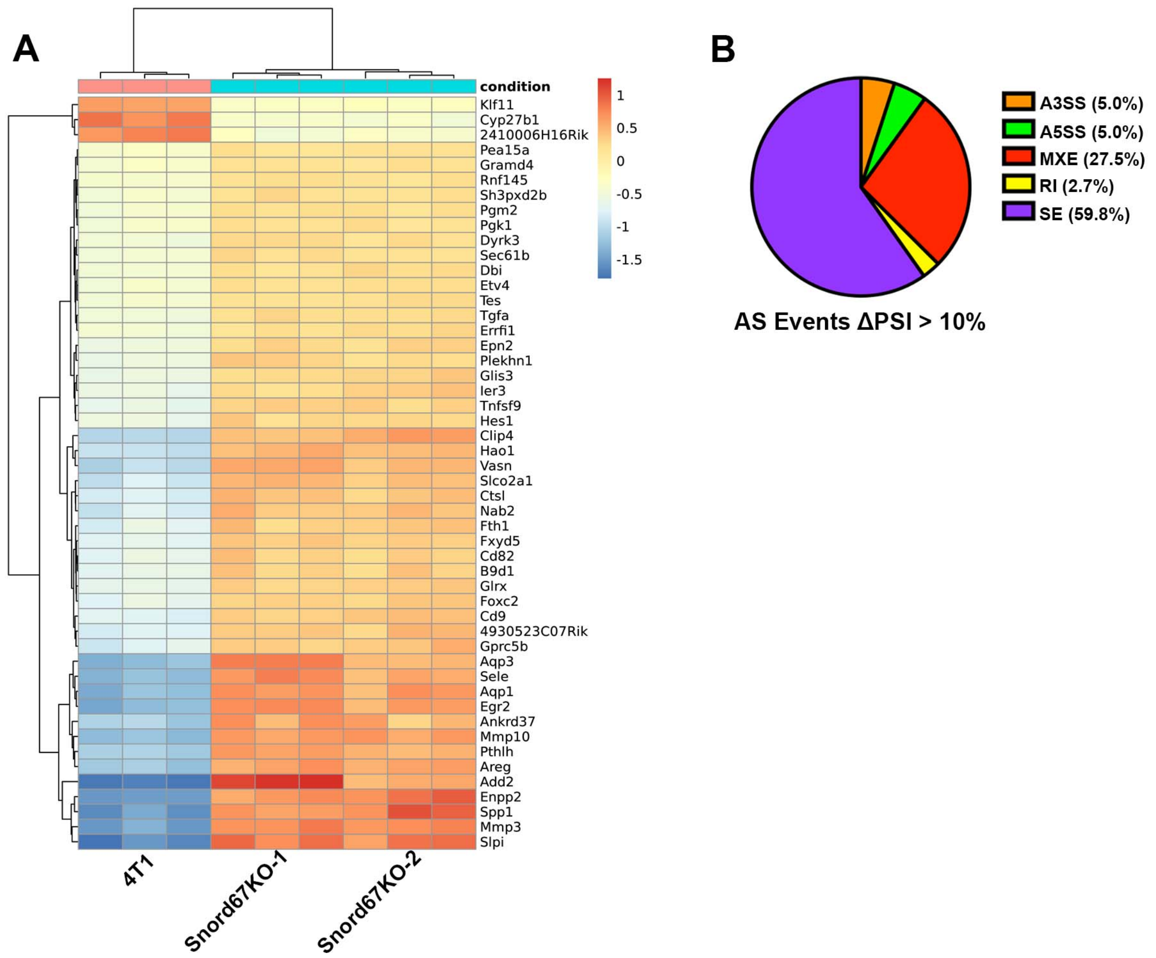


**Supplemental Figure 3. RNA-Seq of 4T1 WT and Snord67 knockout reveals differentially expressed genes.** (**A**) RNA-Seq analysis was mapped using the mouse mm10 genome and visualized as a heatmap showing the top 50 differentially expressed genes with the lowest adjusted p-value/FDR from DESeq2. (**B**) Pie chart showing the distribution for each type of AS event found to be differentially spliced between 4T1 WT and Snord67 knockout cells with ΔPSI>10%.

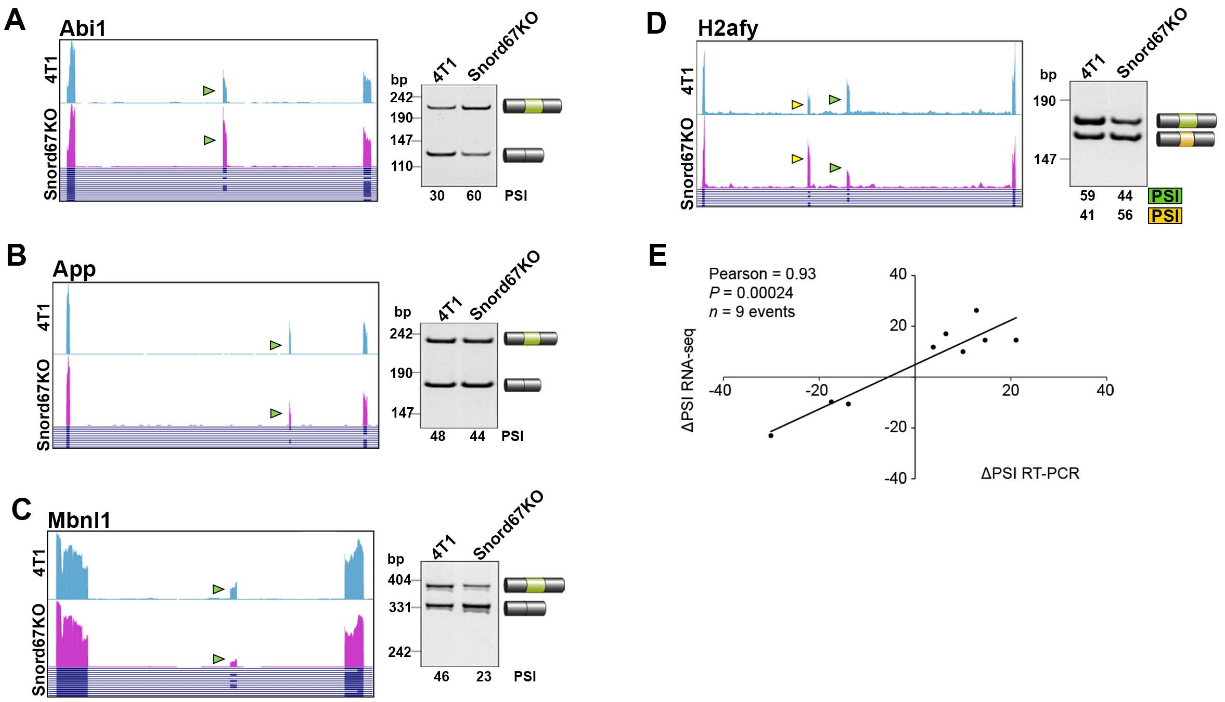


**Supplemental Figure 4. Validation of AS events identified by RNA-Seq in Snord67 knockout compared to 4T1 WT cells.** (**A-D**) Validation by RT-PCR of differential AS events between 4T1 WT and Snord67 knockout cells (Snord67KO). RNA-seq data as displayed by the UCSC browser - mm10 (left) and RT-PCR assays (right) for differentially spliced genes. Arrows indicate alternatively spliced exons. (**E**) Correlation between RNA-sequencing and RT-PCR ΔPSI values for nine selected genes is shown. P was calculated by Student T distribution.

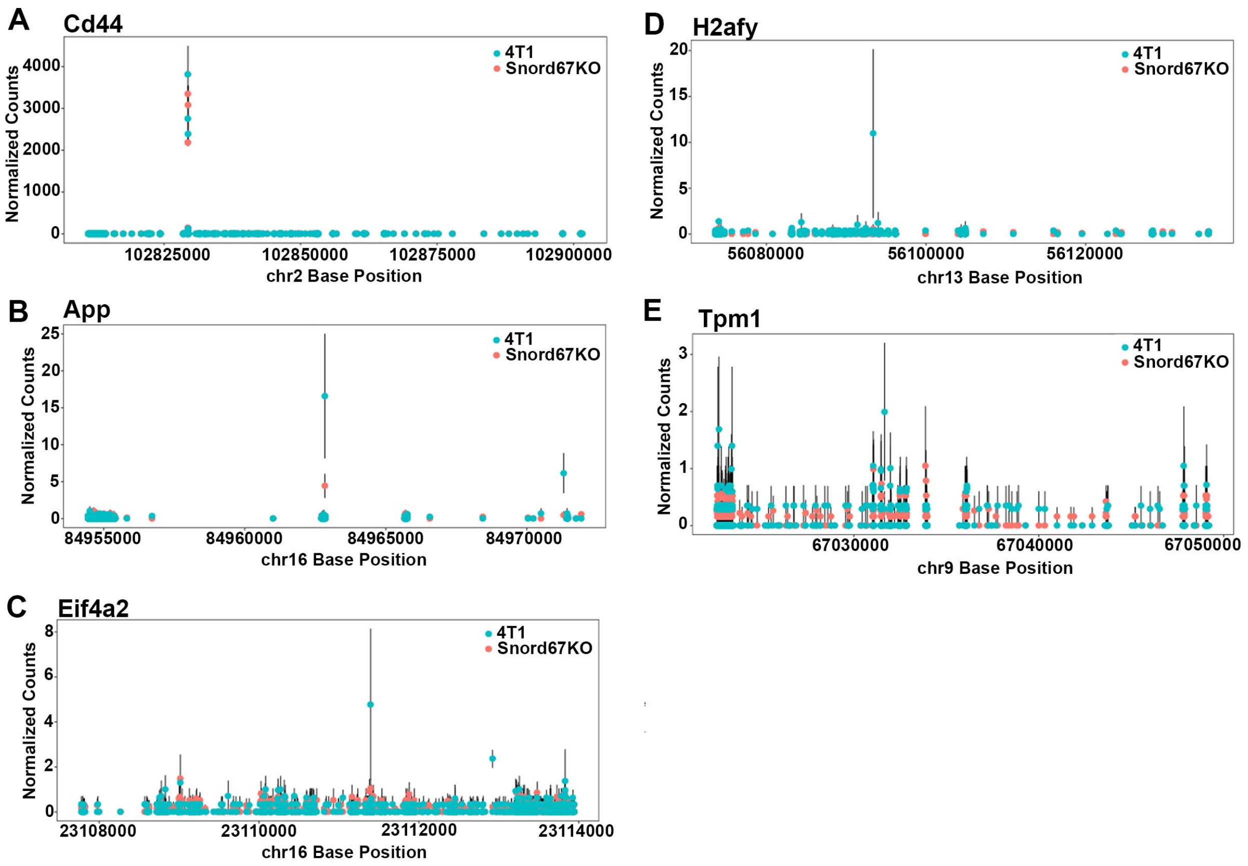


**Supplemental Figure 5. Analysis of 2'-*O*-methylation of pre-mRNA of AS genes shows no difference between 4T1 WT and Snord67 knockouts.** (**A-E**) Methylation mapping by RibOxi-Seq of genes that are differentially spliced between 4T1 and Snord67KOs; methylated reads are aligned by chromosome base position. bp, basepairs; PSI, percent spliced in. Positions with potential differences in methylation underwent secondary analysis, and all were found to be false positives since none had flanking sequences that correspond to the SNORD67 antisense element.

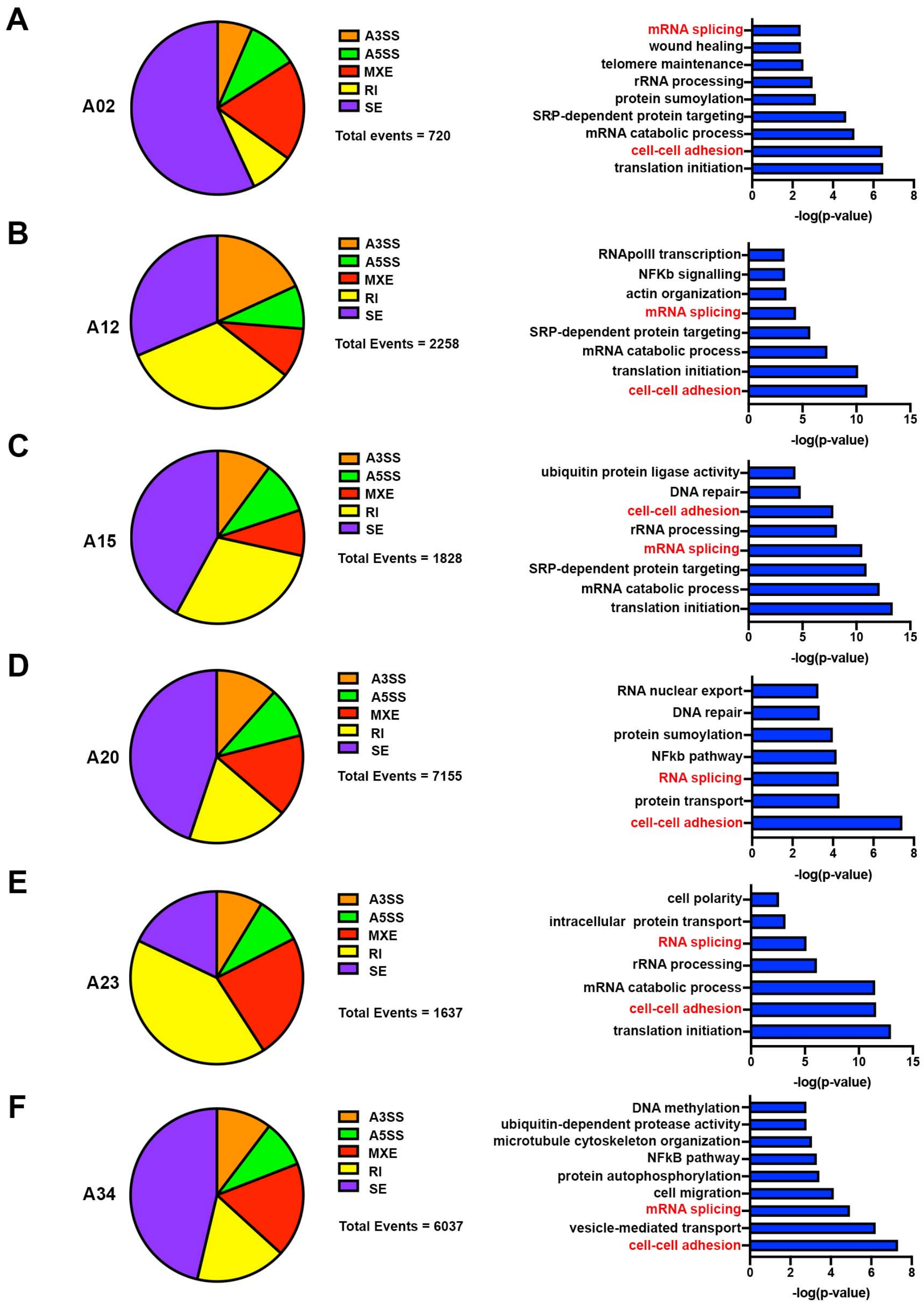


**Supplemental Figure 6. Alternative Splicing events in UNC rapid autopsy program (RAP) samples. a-f**, Pie charts of differential AS event types with Bayes Factor >20 found between primary tumors and LN metastases in breast cancer patients (left panels) and bar charts of gene ontology terms obtained using the DAVID functional annotation tool with p-value<0.05 (right panels). Red text highlights terms that are common between all UNC RAP samples.

| Gene Symbol | Gene Description | Ratio AxLN/ MFP | Ratio AxLN/ Lung Mets |
|---|---|---|---|
| 2410006H16Rik | RIKEN cDNA 2410006H16 gene | 2.44254 | 0.44586 |
| H19 | H19, imprinted maternally expressed transcript | 2.42053 | 0.19068 |
| Neat1 | nuclear paraspeckle assembly transcript 1 (non-protein coding) | 2.40555 | 0.40762 |
| 5430416N02Rik | RIKEN cDNA 5430416N02 gene | 2.25268 | 0.52391 |
| 5430416N02Rik | RIKEN cDNA 5430416N02 gene | 2.11608 | 0.52848 |
| Snord90 | small nucleolar RNA, C/D box 90 | 2.10628 | 0.47847 |
| Snord61 | small nucleolar RNA, C/D box 61 | 2.07915 | 0.47321 |
| Gm26079 | predicted gene, 26079 | 1.93116 | 0.58682 |
| Gm22422 | predicted gene, 22422 | 1.80781 | 0.57619 |
| Snord111 | small nucleolar RNA, C/D box 111 | 1.73464 | 0.47076 |
| Gm25970 | predicted gene, 25970 | 1.71851 | 0.54023 |
| 2810008D09Rik | RIKEN cDNA 2810008D09 gene | 1.68226 | 0.64855 |
| Snord67 | small nucleolar RNA, C/D box 67 | 1.65565 | 0.56170 |
| Gm22858 | predicted gene, 22858 | 1.61493 | 0.57474 |
| Scarna9 | small Cajal body-specific RNA 9 | 1.59294 | 0.60752 |
| Scarna17 | small Cajal body-specific RNA 17 | 1.56472 | 0.68108 |
| Snord1c | small nucleolar RNA, C/D box 1C | 1.54923 | 0.63556 |
| Snord1c | small nucleolar RNA, C/D box 1C | 1.54840 | 0.64509 |
| Gm25835 | predicted gene, 25835 | 1.54222 | 0.63436 |
| Snord104 | small nucleolar RNA, C/D box 104 | 1.53706 | 0.67291 |
| Snora28 | small nucleolar RNA, H/ACA box 28 | 1.51386 | 0.65564 |
| Snord1b | small nucleolar RNA, C/D box 1B | 1.49123 | 0.64902 |
| Gm23119 | predicted gene, 23119 | 1.48750 | 0.71018 |
| Snora34 | small nucleolar RNA, H/ACA box 34 | 1.44901 | 0.68701 |
| Snora16a | small nucleolar RNA, H/ACA box 16A | 1.40703 | 0.67232 |
| Il10rb | interleukin 10 receptor, beta | 1.40046 | 0.64862 |
| Snord16a | small nucleolar RNA, C/D box 16A | 1.34579 | 0.71307 |
| 4930455G09Rik | RIKEN cDNA 4930455G09 gene | 1.24982 | 0.32465 |
| Gm22009 | predicted gene, 22009 | 1.20591 | 0.75911 |
| Gm20412 | predicted gene 20412 | 0.94989 | 0.38742 |

**Supplemental Table 1. Differentially expressed ncRNAs in AxLN tumors compared to MFP tumors and lung metastases**

| | |
|---|---|
| **Primers for DNA Sequencing** | |
| Snord67 F | GCAGTGAAATTTCTTCAGTGTCTG |
| Snord67 R | TGAGACAATGGTGTTATAAGTGTTC |
| **Primers for qPCR** | |
| Universal Stem-Loop qPCR Reverse Primer | TCCCGACCACCACAGCC |
| Snord67 Stem-Loop RT | GCGTGGTCCCGACCACCACAGCCGCCACG ACCACGCTGAGTCAG |
| Snord111 Stem-Loop RT | GCGTGGTCCCGACCACCACAGCCGCCACG ACCACGCATCAGATC |
| Ckap5 F | GGCCTTCGGGTGATTGAGAT |
| Ckap5 R | TGACCTCCATCTGAGGGGAA |
| Sf3b3 F | TACCGAAGCCATCCTTCTTGC |
| Sf3b3 R | CCCGCTGTAAGGTCAGGTTG |
| Gapdh F | AGTATGACTCCACTCACGGCAA |
| Gapdh R | TCTCGCTCCTGGAAGATGGT |
| Rplp0 F | ATCCCTGACGCACCGTGA |
| Rplp0 R | TGCATCTGCTTGGAGCCCACGTT |
| Snord90 F | ATTTCATAGGGCAGATTCTGAG |
| Snord90 R | CTTCAGATTTCATAATAGGACAATCAAT |
| Snord61 F | ATTTGAATCCACTGATCTTCCG |
| Snord61 R | CAAGCTCAGAACTTCTTAGAGGAC |
| Snord111 F | ATTTAATTCATGTCTCTTCTCTGACAT |
| Snord111 R | ATCAGATCAATAAGGCAAAAATCAT |
| Snord104 F | CGGCGATGATGACACTCC |
| Snord104 R | GGCTCAGATTACGATTCGC |
| Snord1C F | GTTGAGCTGAGGATGATTTAAGGTTA |
| Snord1C R | GTCGAGCCTCAGTAAACCATG |
| Snord67 F | TGAGTTGCACACTGGTGGAG |
| Snord67 R | TGAGTCAGATGGCCCCTG |
| **Sequences for Cas9-sgRNA constructs** | |
| Snord67 sgRNA 1 | GCCTGTATCACCTGATACTA TGG |
| Snord67 sgRNA 2 | GGTGACAAAATCAAGTGCAC AGG |
| Snord67 sgRNA 3 | GGCACCACTCAGTACTACCC TGG |
| Snord67 sgRNA 4 | GTGACAAAATCAAGTGCACA GGG |
| Snord111 sgRNA 1 | TTTGCCTTATTGATCTGATC AGG |
| Snord111 sgRNA 2 | AAGGCAAAAATCATCTCCAG AGG |
| Snord111 sgRNA 3 | GAGAAGAGACATGAATTAAA TGG |
| Snord111 sgRNA 4 | TCTTCTCTGACATTCTCCTC TGG |
| **Sequences for ASOs** | |
| Snord67 ASO | ATGGCCCCTGTGCACTTGAT |
| NC ASO | AACACGTCTATACGC |
| **Primers for alternative splicing RT-PCR** | |
| Abi1 F | CAATTCTCTGCTCAGCCTCA |
| Abi1 R | GGCGGAGAGTCATCAAACAT |
| Mbnl1 F | TGACTGTCGGTTTGCTCATC |
| Mbnl1 R | GCATGTTGGCTAGAGCCTGT |
| Setd3 F | GCTCTGGTTGAGAAAATACGG |
| Setd3 R | CTGAATTCTTAGCAGATTCAACAG |

| | |
|---|---|
| Stk25 F | GACCGATATAAGCGCTGGAA |
| Stk25 R | CGGACTAGTGTGGACAAGCA |
| Zfp68 F | CTCATCTGCCTCTTCAACC |
| Zfp68 F | TCCAGCATCACATCCCTGTA |
| Cd44 F | TCTTTATCCGGAGCACCTTG |
| Cd44 R | TGGGTCGAAGAAATCTGTCC |
| App F | ATTCTTTTACGGCGGATGTG |
| App R | CTTTGGCTTTCTGGAAATGG |
| Eif4a2 F | GGATTGACGTGCAACAAGTG |
| Eif4a2 R | CCAAATCGACCCCCTCTG |
| H2afy F | GACGGCTTCACTGTCCTCTC |
| H2afy R | GCCCTTCTTCTCCAGTGTGT |
| Tpm1 F | GTGGCCCGTAAGCTGGTC |
| Tpm1 R | TCATATTTGTCTTCCTTCTGAGAGT |

**Supplemental Table 2. Sequences for Primers, CRISPR sgRNAs, and ASOs**

| | |
|---|---|
| NEB miRNA linker | 5´-/rApp/CTGTAGGCACCATCAAT/NH2/- 3´ |
| RibOxi RT Primer | 5'-GTGACTGGAGTTCAGACGTGTGCTCTTCCGATCTNNNNNNATTGATGGTGCCTACAG-3' |
| 5'RNA linker | 5'-/Biosg/ACACUCUUUCCCUACACGACGCUCUUCCGAUCUNNNN-3' |
| PCR_i5 | 5'-aatgatacggcgaccaccgagatctacac**-i5(8nt)-**acactctttccctacacgacgctcttccgatct-3' |
| PCR_i7 | 5'-caagcagaagacggcatacgagat**-i7(8nt)-**gtgactggagttcagacgtgtgctcttccgatct-3' |

**Supplemental Table 3. Primers and Oligos used for RNA- and RibOxi-Sequencing**
